# Striatal dopamine modulates reward-reinforced temporal learning in humans

**DOI:** 10.1101/2025.07.03.662974

**Authors:** Emily K. DiMarco, Angela Jiang, Ashley R. Shipp, Stephen B. Tatter, Adrian W. Laxton, Kenneth T. Kishida

## Abstract

The neurobiological mechanisms underlying human time perception remain elusive. Evidence has consistently linked striatal dopamine to timing behaviors, but it is still uncertain how rapid changes in dopamine may modulate human time perception. Many tasks designed to measure time perception utilize instrumental conditioning paradigms that reinforce correctly timed intervals. In these tasks, subjects are shown to improve their performance following repeated presentations of temporal cues – a phenomenon known as ‘temporal learning’. We sought to determine the association between rapid changes in human dopamine levels and temporal learning on an interval timing task that tested the reproduction of 1000ms, 3000ms, and 5000ms intervals in the presence and absence of monetary reinforcement. We utilized human voltammetry to measure real-time dopamine concentrations from the striatum of patients with Parkinson’s disease while they performed the interval timing task. We first compared task behavior between patients with Parkinson’s disease and neurologically healthy controls and found significant differences in the reproduction of 1000ms intervals, but not 3000ms or 5000ms intervals of time. Further, we observed that during 1000ms intervals, increases in striatal dopamine concentrations were associated with increases in temporal errors, but only during the expectation of monetary reinforcement. We also demonstrated that as temporal errors decrease overtime during temporal learning, so do striatal dopamine concentrations. These results suggest that dopamine may be driving temporal learning through the generation of temporal errors in response to positive reinforcement. These findings may have significant implications in our understanding of the role that dopamine plays in time perception.

**Significance Statement:** Human time perception is a fundamental cognitive ability but the interaction between how the human brain perceives time and dopamine’s role is unclear. This study is the first of its kind to apply human voltammetry to measure rapid changes in dopamine levels associated with temporal learning in the presence and absence of positive reinforcement. We demonstrate that temporal learning may be affected by moment-to-moment changes in dopamine levels, which are also counterintuitively related to the generation of temporal errors in the presence of expectations of rewarding feedback.

## Introduction

Timing and timed behaviors are essential to how one learns from and navigates the world; however, how humans perceive time has long been an open question. Research has consistently implicated striatal dopamine in the modulation of timing in mammals, including humans (Meck 1996; Fung et al. 2021). However, it remains unclear how task structure, particularly tasks that utilize rewards to train subjects, relate to dopamine’s role in timing. Therefore, investigating how dopamine is associated with timing in the presence and absence of reinforcement may provide a clearer picture of dopamine’s role in processes underlying how the human perceives time.

While many task designs have been employed to measure timing behaviors, peak interval procedures are one common example of a task used to investigate effects of dopamine on the reproduction of time intervals in the presence of reinforcement (MacInnis & Guilhardi, 2006). Peak interval procedures typically begin with the presentation of a temporal cue to “start the clock” – like the dropping of a lever in rodent studies – followed by some action to reproduce the duration of time cued to “stop the clock” – like a lever press. This behavior is then reinforced with a reward after the cued duration has elapsed. In these studies, subjects are often trained to achieve a certain threshold of correctness after repeated exposure to the temporal cues (Rakitin et al., 1998).

As task structures like peak interval procedures, and many others employed to investigate time perception, utilize instrumental conditioning to train and measure timing, it is imperative to better understand the role that predictable cues and reinforcement play in timing behaviors (Fung et al. 2021). Previous work utilizing peak interval procedures has shown that subjects generate and repeatedly update “expected values” for each temporal cue by weighing the cumulative values of past reinforcements against the current temporal cue’s reinforcement value (DiMarco et al., 2024). This work additionally shows that subjects yield an initial increase in temporal errors, followed by a continuous reduction in temporal errors with repeated exposure to temporal cues, in a process known as temporal learning (DiMarco et al., 2024). Because instrumental conditioning tasks are commonly used to investigate the role of dopamine in reinforcement learning processes (Kishida & Sands, 2021; Sands et al., 2024), and substantial past research has queried how rewards and timing intersect (Hollerman & Shultz, 1998; Meck, 2014; Fung et al., 2021), understanding dopamine’s role in tasks that utilize instrumental conditioning to measure timing behavior may help clarify dopamine’s role in time perception.

In this study, we designed a peak interval procedure that tests the reproduction of 1000ms, 3000ms, and 5000ms cues in the presence and absence of reinforcement. Based on previous work (DiMarco et al. 2024), we hypothesized that increased striatal dopamine concentrations would be associated with increases in temporal errors in the presence, but not the absence, of reinforcement for short (e.g., 1000ms) intervals. We collected and compared performance on this task from patients with Parkinson’s disease and neurologically healthy controls. In patients with Parkinson’s disease, we also measured dopamine levels in the striatum using human voltammetry (Kishida et al., 2011; Kishida et al., 2016; Montague & Kishida, 2018; Bang et al., 2020, Liebenow et al., 2023; Sands et al., 2023; Sadibolova et al., 2024), which provided 100ms temporal resolution of changes in dopamine levels throughout the task. Our results suggest that dopamine availability may modulate the reproduction of 1000ms durations. We did not observe associated effects in 3000ms or 5000ms durations. Further investigation of temporal behavior on 1000ms duration cues demonstrated that reward expectations are associated with an initial increase in temporal error that is overcome throughout subsequent cue presentations in the presence of reinforcement, demonstrating temporal learning. To determine if temporal learning was associated with dopamine levels, we applied human voltammetry (Kishida et al., 2011; Kishida et al., 2016; Montague & Kishida, 2018; Bang et al., 2020, Liebenow et al., 2023; Sands et al., 2023; Sadibolova et al., 2024) to measure real-time striatal dopamine concentrations. We found significant associations between increased dopamine concentrations and temporal errors for reinforced, but not unreinforced, cues during 1000ms intervals. This work may begin to help clarify the role that dopamine plays in human time perception and help to better understand the potential common dopaminergic mechanisms underlying time perception and instrumental conditioning.

## Methods

### Subjects

We recruited and consented 5 patients with Parkinson’s disease from Atrium Health Wake Forest Baptist Medical Center in Winston-Salem, North Carolina using methods approved by the Wake Forest University School of Medicine Institutional Review Board (IRB00017138) (**Table 1**). Recruited patients were scheduled to receive standard-of-care deep brain stimulation surgery (DBS) for the treatment of Parkinson’s disease symptoms. During the same visit, they were consented into the study and performed a behavioral demonstration of the task about a week prior to their standard-of-care DBS surgery. During the DBS electrode implantation surgery, task behavior and voltammetry measurements were collected concurrently for a maximum of thirty minutes. One patient was unable to complete the full task, but their data (up to the point that they decided to stop) was included in our analyses. In addition, n = 40 neurologically healthy (no diagnosis of Parkinson’s disease) participants were recruited from the Winston-Salem, North Carolina region and consented using methods approved by the Wake Forest University School of Medicine IRB (IRB00042265) (**Table 1**). All neurologically healthy controls performed the same timing task as performed by the Parkinson’s disease cohort.

**Table 1.**
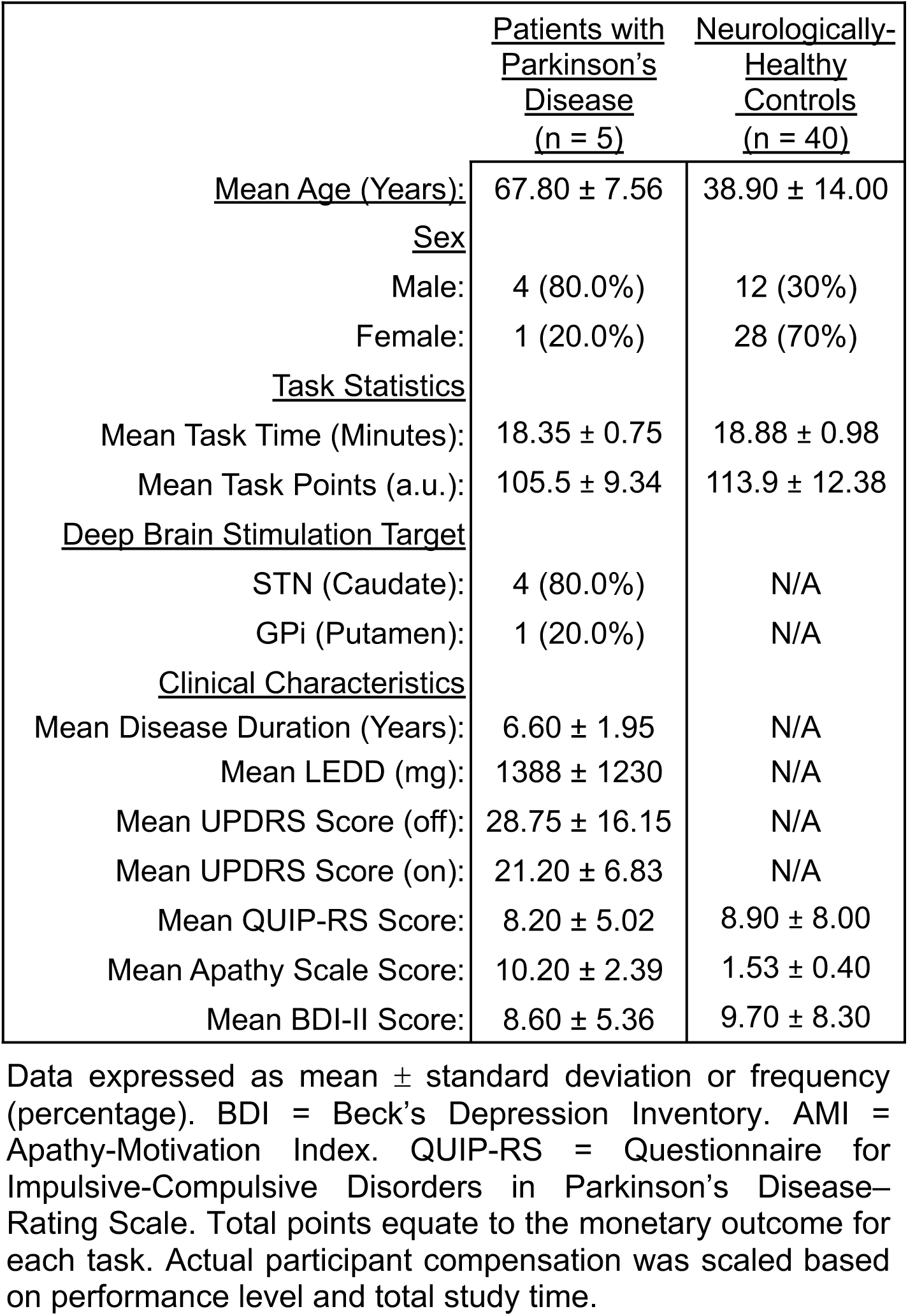
Demographics and task performance

### Task Design

The peak interval procedure used in this study is an interval timing task with 120 trials broken into three phases (**Fig.1A**). For each trial, participants were presented with a demonstrator cue for one of three criterion durations: 1000, 3000, or 5000ms, then asked to reproduce the duration of time with a button press on a button box after a reproduction cue was presented (to signal the start of the reproduction duration). On initialization of the task, 12 visual cues were randomly selected from a pool of 60 fractal images sourced from free online stock images. In phase I (48 trials), six cues were randomly selected from an array and presented 8 times each, two different cues for each criterion duration of 1000ms, 3000ms, and 5000ms (denoted as: 1000ms Neutral -1, 1000ms Neutral-2, 3000ms Neutral-1, 3000ms Neutral-2, 5000ms Neutral-1, and 5000ms Neutral-2). Participants received no feedback in phase I. In phase II (48 trials), six new cues were selected from an array and presented 8 times each. For one of each of the 1000ms, 3000ms, and 5000ms cues, participants were positively reinforced based on their performance (earned up to $3) (denoted as: 1000ms Win, 3000ms Win, and 5000ms Win). For the remaining three cues, no reinforcement was presented (denoted as: 1000ms Neutral, 3000ms Neutral, 5000ms Neutral). In phase III (24 trials), the three reinforced cues from phase II were presented 8 times each, but without reinforcement. The three cues probabilistically cued the other possible criterion durations (CDs) 90% of the time. The other 10% of the time, they cued the original duration presented in phase II (**Fig. 1B**).

**Figure 1.**
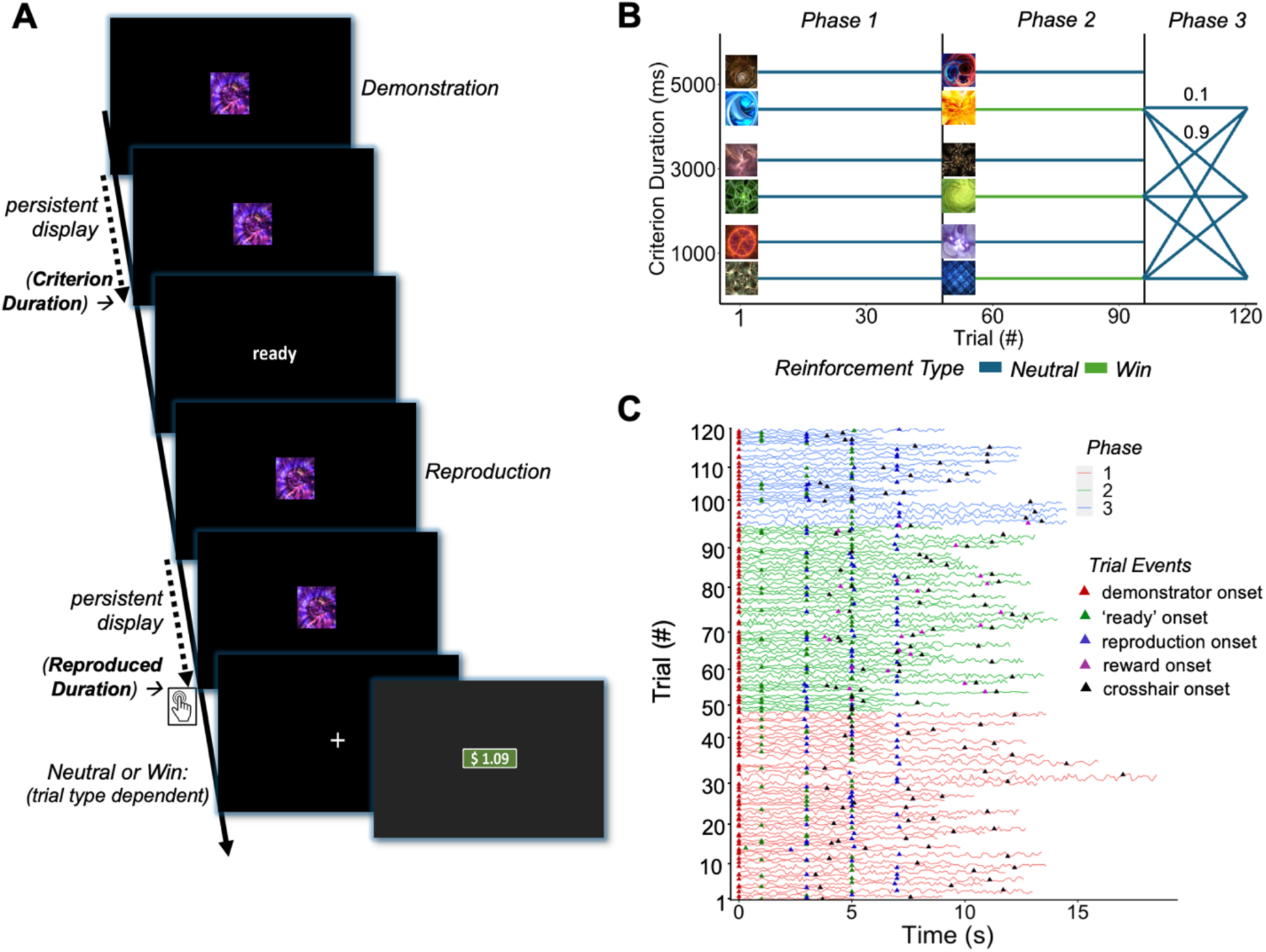
Task schematic and voltammetry timeseries. **A.** The Peak Interval Procedure 3-Phase (PIP-3P) is an interval timing task that has 120 trials broken into three phases. Subjects are presented with a demonstrator cue for either 1000ms, 3000ms, or 5000ms, then asked to reproduce the duration of time with a button press on a button box. **B.** In Phase I (48 trials), six cues are randomly presented 8 times each, two different neutral cues for each 1000ms, 3000ms, and 5000ms. Subjects receive no feedback in Phase I. In Phase II (48 trials), six novel cues are randomly presented 8 times each. For one of the 1000ms, 3000ms, and 5000ms cues in phase II, subjects are positively reinforced with monetary feedback scaled to their performance (win cues; earn up to $3). For the remaining three cues, no reinforcement is presented (neutral cues). In Phase III (24 trials), the three Win cues from Phase II are presented 8 times each, but without reinforcement. The three cues now probabilistically cue the other possible criterion durations (CDs) 90% of the time. The other 10% of the time, they cue the original duration presented in Phase II. **C.** Z-scored dopamine concentrations from human voltammetry measurements for each trial on the PIP-3P taken from Parkinson’s disease patient 1. Each trial is broken down into events of demonstrator cue onset (red triangle), ‘ready’ prompt onset (green triangle), reproduction cue onset (blue triangle), reward onset (purple triangle), and crosshair onset (black triangle).

Stages of the task included: demonstrator cue, ready prompt, reproduction cue, and reinforcement (if presented) (**Fig. 1A**). Upon presentation of the demonstrator cue, participants observed the cue for the assigned duration (1000ms, 3000ms, or 5000ms). Following the offset of the demonstrator cue, a black screen appeared for 500ms before a ‘ready’ prompt was displayed for 1000ms. Following another black screen for 500ms, the reproduction cue appeared, which was the same cue as the demonstrator cue. Subjects were instructed to use the hand-held button box to reproduce the duration of time by pressing the button when the demonstrator cued duration elapsed. The cue disappeared when the button was pressed or after twice the demonstrator cue duration passed. Immediately after the participant pressed the button, a monetary reinforcement screen or a hairpin cross screen was displayed based on the trial type. In between trials, a black screen with a crosshair in the center of the screen was shown for an interval of time randomly drawn from a Poisson distribution with a characteristic mean and variance both equal to 1500ms. On average, experiments lasted ∼18 minutes per participant (**Table 1**).

Following completion of the task, all neurologically healthy participants completed several demographic and clinical questionnaires to better understand and control for the potential individual differences in task performance that have been known to affect timing and reinforcement learning behaviors. These clinical questionnaires included measures of depression (Beck’s Depression Inventory-II), apathy (Apathy-Motivation Index), and impulsivity (Questionnaire for Impulsive and Compulsive Disorders in Parkinson’s Disease-Rating Scale). At the end of the study visit, participants were compensated based on the study duration ($5 per 15 minutes), with a bonus based on task performance (ranging from $20 - $60 additional). The task was coded using the PyGame toolbox in Python.

### Human Fast-Scan Cyclic Voltammetry (FSCV)

Patients with Parkinson’s disease who elected to undergo deep brain stimulation surgery for the treatment of the motor symptoms of their disorder were consented into the study. During their standard-of-care neurosurgery, fast scan cyclic voltammetry (FSCV) was performed using carbon fiber microelectrodes surgically placed in the caudate (4 patients) or putamen (1 patient) as determined by neurosurgical planning prior to the surgeries. Concurrently with the collected FSCV measurements, participants performed the described peak interval procedure. Behavior and subsecond dopamine concentrations were then compared for each patient (**Fig. 1C**).

The human FSCV protocol applied in the current study is described in previous publications (Kishida et al. 2011; Kishida et al. 2016; Montague & Kishida 2018; Bang et al. 2020, Liebenow et al. 2023; Sands et al. 2023; Sadibolova et al. 2024). Briefly, after the carbon fiber microelectrode was inserted along the planned DBS surgical trajectory the FSCV protocol was applied. The FSCV protocol consisted of an equilibration phase and an experiment phase. During the equilibration phase, a triangular waveform was applied, ramping up from a holding potential of -0.6 V to a peak of +1.4 V at 400 V/s and ramping back down to -0.6 V at -400 V/s, and repeating at 60 Hz. This phase provided time for the equilibration of the electrode surface (∼10 minutes). Following equilibration of the electrode, the experimental phase began, and the same waveform was applied, but cycled at 10 Hz for the duration of the experimental window (∼18 minutes).

Dopamine concentrations were derived using a generalized linear regression model that had been trained on flow-cell data, consistent with previous protocols (Kishida et al., 2016). Data to train the models was collected using 8 electrodes under the same FSCV protocols described previously (Kishida et al. 2011; Kishida et al. 2016; Montague & Kishida 2018; Bang et al. 2020, Liebenow et al. 2023; Sands et al. 2023; Sadibolova et al. 2024). Each electrode was exposed to solutions of varying concentrations (0 - 4000 nM) of dopamine diluted in phosphate-buffered saline while recording electrochemical data at 10 Hz. Overall, 60 solutions were used in the training set, with 120 seconds of data from each solution. Data during movement artifacts caused by injecting solutions into the flow cell were excluded from model training leaving 15,600 voltammograms to train the model.

Current (nA) measurements within each voltammogram were sampled at a rate of 250kHz, which resulted in 2500 samples in each 10-millisecond triangular waveform. To reduce the effect of electrical noise on measurements, voltammograms were down sampled from 2500 samples to 42 samples by averaging every 60-sample bin. An elastic-net regularized linear regression model was trained on these down sampled voltammograms (42 samples) concatenated with the first (41 samples) and second derivative (40 samples) of the voltammograms for a total of 123 independent variables (𝑥) to predict the concentration of dopamine:

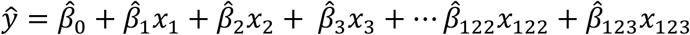

where 𝑦̂ is the estimated concentration of dopamine, 𝛽̂_0_ is the intercept, and 𝛽̂_1_ to 𝛽̂_123_ are the coefficients for each independent variable. Here, 𝑥_1_ to 𝑥_42_ are current values for each point in the down sampled voltammogram, 𝑥_43_ to 𝑥_83_ are values for points in the 1st derivative, and 𝑥_84_ to 𝑥_123_ values for points in the second derivative. The elastic net mixing parameter (𝛼) was set to 1 for lasso regularization, which emphasizes variable selection by forcing some coefficients to be 0, effectively performing feature selection. Model training was performed using the glmnet package in MATLAB 2019a (Qian, et al., 2013). The resulting model was validated on out-of-sample testing data from a single electrode held out from training, yielding an r^2^ of 0.95 between normalized predicted dopamine concentrations and their corresponding known values.

### Statistical Analyses

Analyses for the human voltammetry experiments and behavior were completed using a combination of R Studio (RStudio Team, 2021) and Matlab R2022b (The MathWorks Inc., 2022). For the behavior from the peak interval procedure, absolute temporal error was calculated using the absolute value of the difference between the criterion duration (1000ms, 3000ms, or 5000ms) and the reproduced duration of the subject. Learning curves were calculated by first separating the data into groups based on cue type (i.e. 1000ms Win), and then plotting the absolute temporal error over all eight cue appearances for each subject. Curves were individually fit to the mean absolute temporal error of each subject at each appearance using an exponential function:

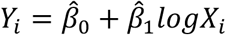

where Y is absolute temporal error for each cue (i). X is each appearance of the cue (i). 𝛽̂_0_ represents the intercept and measures the value where the line crosses the y-axis. 𝛽̂_1_ is the slope of the curve and measures the rate of change in absolute error, or the learning rate of each subject.

For the human voltammetry experiments, dopamine concentrations (nm) were z-scored patient-wise (5) for each cue type (12). Mean z-scored dopamine concentrations for each cue type were then used in linear regression models to compare dopamine’s relationship with accuracy error and number of cue appearances (i.e. y = mx + b, where y = z-scored dopamine and x = cue appearance number). Following removal of any missed trials, linear regression models were calculated first for each patient, then the group mean.

## Results

### Timing in Parkinson’s disease versus neurologically healthy controls

To begin, timing behavior between patients with Parkinson’s disease (pd) and neurologically healthy controls (healthy) on the peak interval procedure was compared. Two-sample t-tests revealed significant group differences in the reproduction of 1000ms intervals of time for the phase I 1000ms Neutral-1 (**Fig. 2A**; t(43) = -2.3357, p = 0.02424*) and 1000ms Neutral-2 (**Fig. 2B**; t(43) = -2.0195, p = 0.04969*) cues and the phase II 1000ms Neutral (**Fig. 2C**; t(43) = -2.8266, p = 0.007111**) and 1000ms Win (**Fig. 2D**; t(43) = -2.1923, p = 0.03382*) cues. Two-sample t-tests comparing the reproduction of 3000ms and 5000ms intervals showed no significant differences between the reproduced durations of the two groups (phase I cues - 3000 Neutral-1: t(43) =-0.94962, p = 0.3924; 3000 Neutral-2: t(43) =1.2523, p = 0.2766; 5000 Neutral-1: t(43) = 1.7156, p = 0.1414; 5000 Neutral-2: t(43) = -0.2106, p = 0.8430; phase II cues - 3000 Win: t(43) = -0.6062, p = 0.5746; 3000 Neutral: t(43) = -0.5658, p = 0.6004; 5000 Win: t(43) = 0.0035, p = 0.9973; 5000 neutral: t(43) = 2.2873, p = 0.0599).

**Figure 2.**
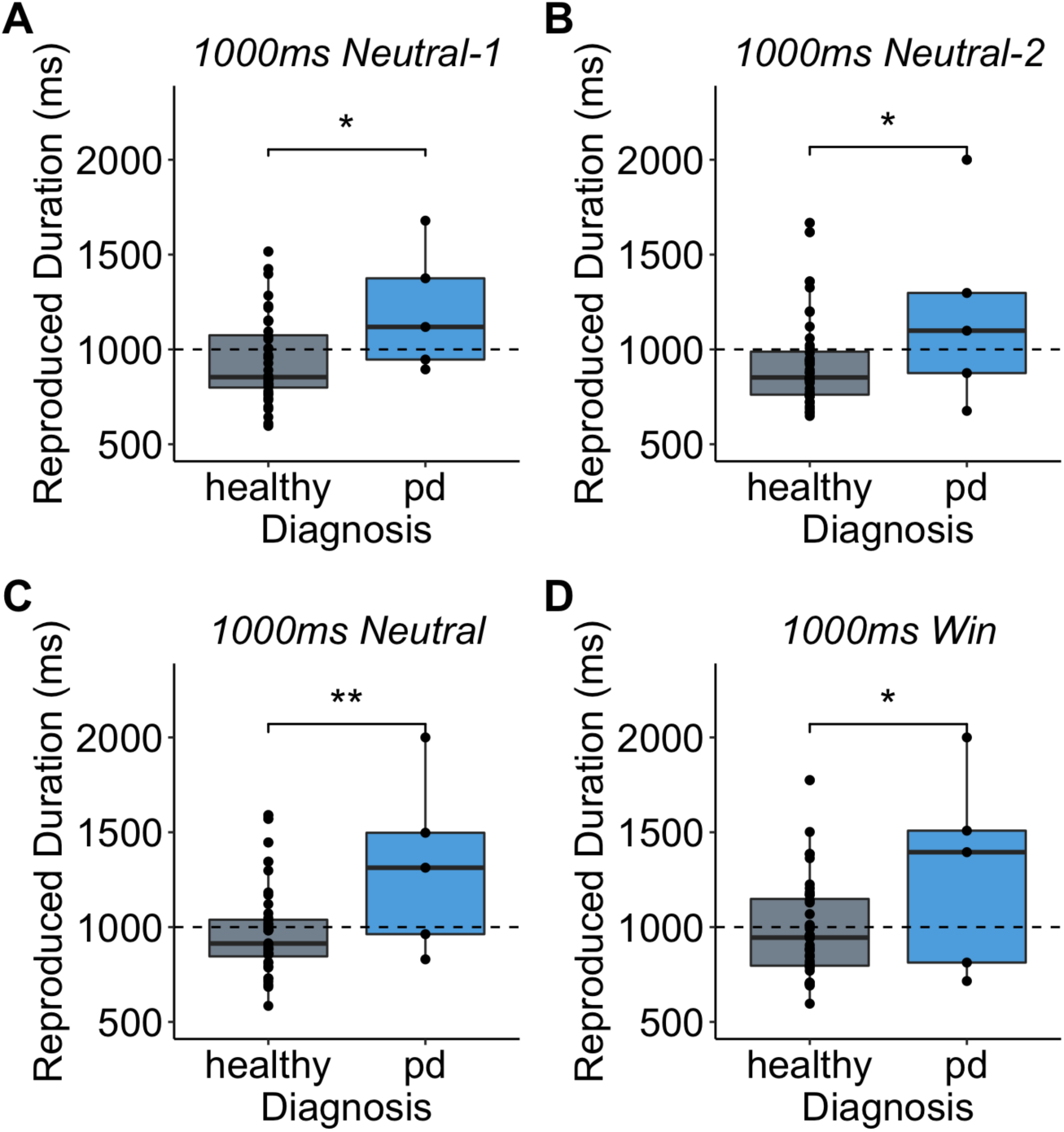
Mean reproduced durations. **A.** for the phase I 1000ms Neutral-1 cue, **B.** for the phase I 1000ms Neutral-2 cue, **C.** for the phase II 1000ms Neutral cue, and **D.** for the phase II 1000ms Win cue. Healthy represents the neurologically healthy controls (40) and pd represents the patients with Parkinson’s disease. Each point denotes the mean reproduced duration of a subject for that cue type. Comparisons based on two-sample t-tests. Significance based on p < 0.05*, 0.01**.

Temporal learning was quantified by calculating the absolute temporal error (ms) for each repeated appearance of cues of the same type. Healthy subjects showed significant temporal learning for the 1000ms duration cues, as paired two sample t-tests comparing absolute temporal error at the first appearance of each cue and the last appearance of each cue revealed significant differences between groups (**Fig. 3**; 1000ms Neutral-1: t(39) = 2.5283, p = 0.0156*; 1000ms Neutral-2: t(39) = 1.5306, p = 0.1339; 1000ms Neutral: t(39) = 1.1692, p = 0.2494; 1000ms Win: t(39) = 2.6077, p = 0.0129*). Based on a paired two sample t-test, healthy subjects also exhibited a significant increase in absolute temporal error between the last appearance of the 1000ms cues in phase I and the first appearance of novel 1000ms cues in phase II (t(79) = -2.5873, p = 0.0115*). Patients with Parkinson’s disease did not exhibit significant temporal learning, as paired two sample t-tests showed no significant differences between the first and last appearances of the 1000ms cues (1000ms Neutral-1: t(4) = -0.8968, p = 0.4205; 1000ms Neutral-2: t(4) = 0.4251, p = 0.6926; 1000ms Neutral: t(4) = 0.7139, p = 0.4995; 1000ms Win: t(4) = 1.5400, p = 0.1874).

**Figure 3.**
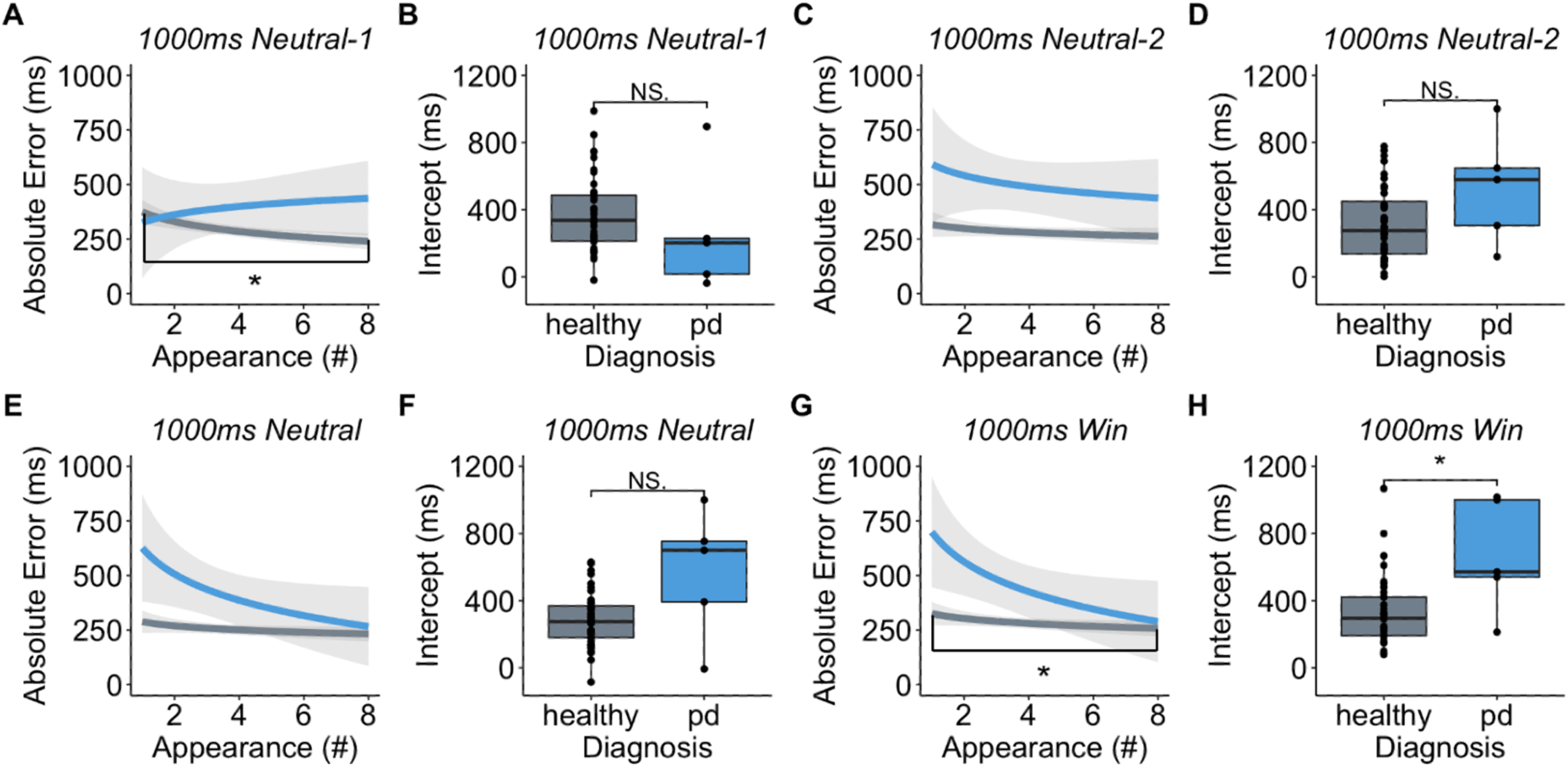
Mean learning curves and intercept values A.B. for the phase I 1000ms Neutral-1 cue, **C.D.** for the phase I 1000ms Neutral-2 cue, **E.F.** for the phase II 1000ms Neutral cue, and **G.H.** for the phase II 1000ms Win cue. Learning curves fit with an exponential function. Intercept derived from the beta value of that function. Comparisons based on Wilcoxon rank sum exact tests. Significance based on p < 0.05*.

To compare temporal learning between patients with Parkinson’s disease and healthy controls, slopes and intercepts of learning curves were calculated and compared (**Fig. 3**). Wilcoxon rank sum exact tests comparing the mean intercept values between patients with Parkinson’s disease and healthy controls showed significant differences for the 1000ms Win cue type (**Fig. 3H**; W = 37, p-value = 0.0205*) and trending differences for the 1000ms Neutral cue type (**Fig. 3F**; W = 47, p-value = 0.05622), but no significant differences for the 1000ms Neutral-1 (**Fig. 3B**; W = 139, p-value = 0.1696) or 1000ms Neutral-2 (**Fig. 3D**; W = 57, p-value = 0.1277) cue types. In patients with Parkinson’s disease, one-way ANOVAs were used to compare the slope and intercept values of the learning curves between each cue type. Slope (F(3,9) = 1.259, p = 0.3465) and intercept (F(3,9) = 0.99749, p = 0.4378) values were not significantly different between cue types in patients with Parkinson’s disease (**Fig. 3A**: intercept = 260.9570, slope = 29.9929; **Fig. 3C**: intercept = 530.4571, slope = -8.2238; **Fig. 3E**: intercept = 567.8429, slope = -19.1595; **Fig. 3G**: intercept = 668.3571, slope = -36.4071).

### Dopamine in temporal learning

To determine the role that dopamine plays in temporal learning, mean z-scored dopamine concentrations of each cue type for each patient were calculated and their associations with temporal learning measures were explored. Linear regression models were used to determine if the absolute temporal error of each patient for each cue type was associated with mean dopamine concentrations for the same cue types (**Fig. 4**). For the 1000ms Neutral-1 (**Fig. 4A**; F(1,30) = 0.5965, p = 0.4460, R^2^ = 0.0195), 1000ms Neutral-2 (**Fig. 4B**; F(1,27) = 1.2340, p = 0.2764, R^2^ = 0.0437), and 1000ms Neutral cues (**Fig. 4C**; F(1,26) = 0.4717, p = 0.4983, R^2^ = 0.0178), no association between absolute temporal error and dopamine concentration was found. For the 1000ms Win cue, absolute temporal error was significantly associated with dopamine concentration (**Fig. 4D**; F(1,25) = 4.596, p = 0.0420*, R^2^ = 0.1553).

**Figure 4.**
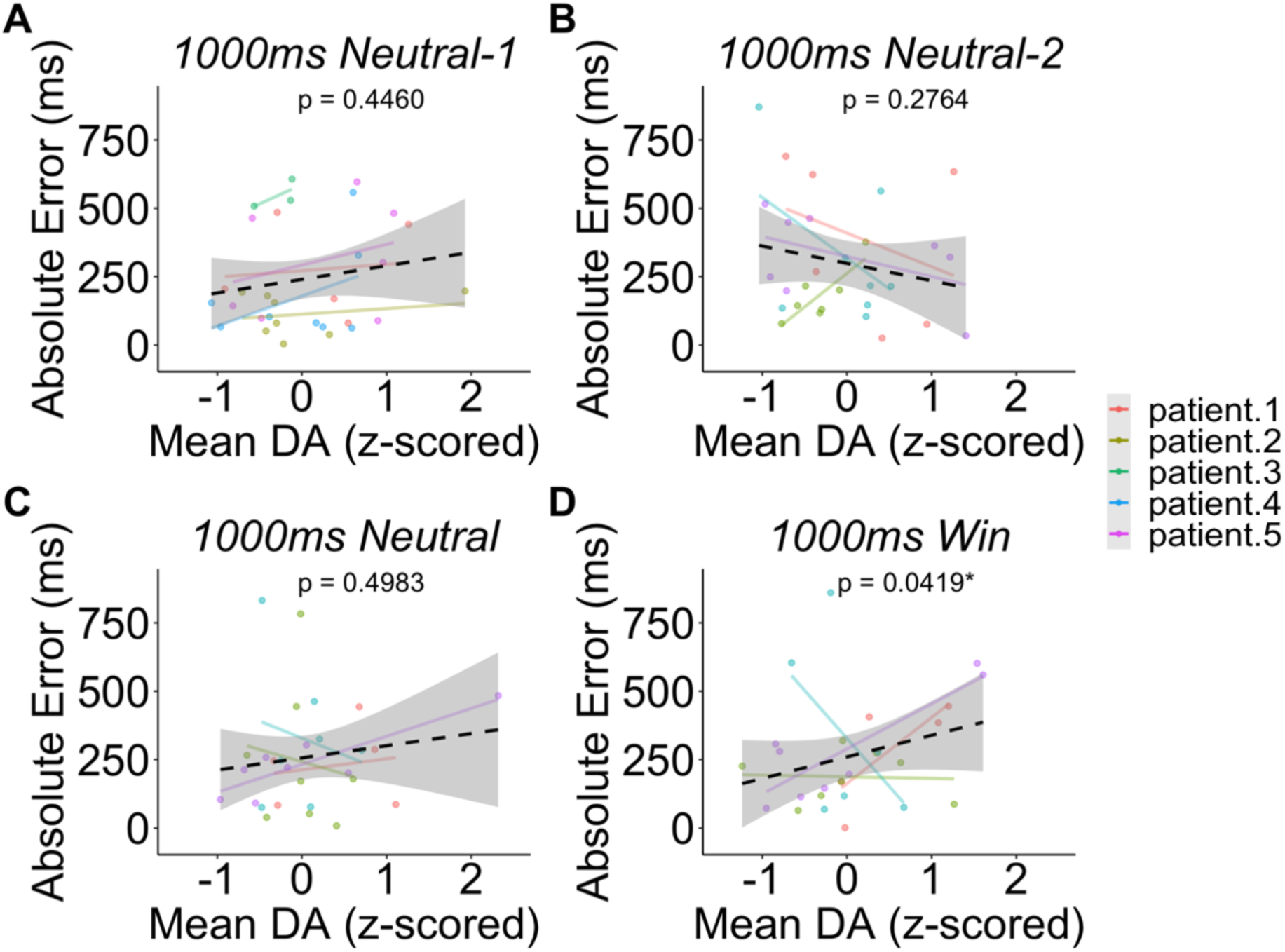
Associations between mean dopamine (DA) concentrations and absolute temporal error. **A.** for the phase I 1000ms Neutral-1 cue, **B.** for the phase I 1000ms Neutral-2 cue, **C.** for the phase II 1000ms Neutral cue, and **D.** for the phase II 1000ms Win cue. Colored lines and points denote individual patient associations. Dotted black line denotes mean association. Associations based on linear regression models. Significance based on p < 0.05*.

Linear regression models were also used to determine the association between dopamine concentrations and cue appearance (**Fig. 5**). For the 1000ms Neutral-1 (**Fig. 5A**; F(1,38) = 0.1978, p = 0.6590, R^2^ = 0.0052), 1000ms Neutral-2 (**Fig. 5B**; F(1,38) = 1.639, p = 0.2083, R^2^ = 0.0413), and 1000 Neutral cues (**Fig. 5C**; F(1,33) = 0.109, p = 0.7433, R^2^ = 0.0033), there were no associations between cue appearances and dopamine concentrations. However, for the 1000ms Win cue, appearance number was significantly associated with dopamine concentration (**Fig. 5D**; F(1,34) = 4.724, p = 0.0368*, R^2^ = 0.1220).

**Figure 5.**
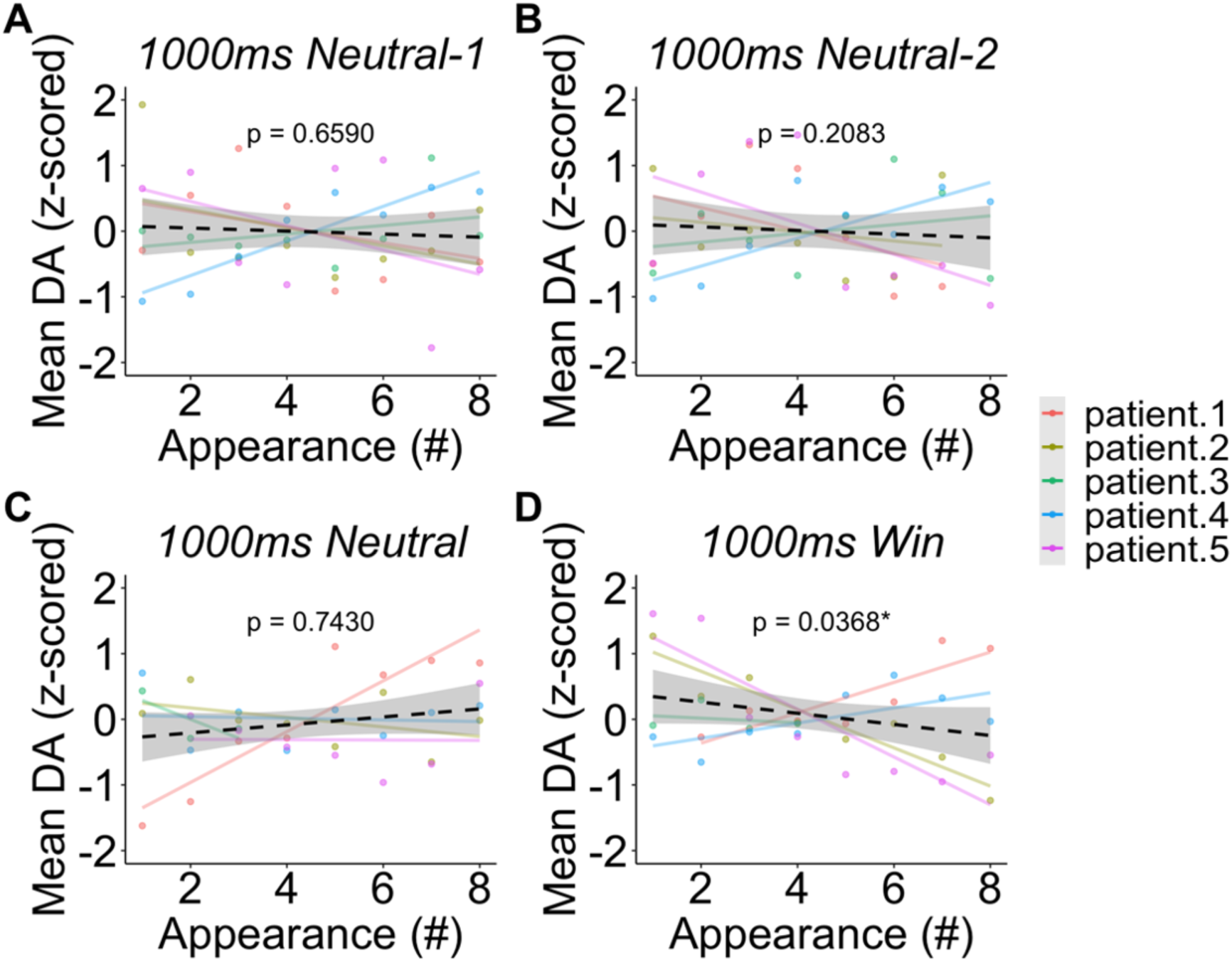
Associations between cue appearance and mean dopamine (DA) concentrations. **A.** for the phase I 1000ms Neutral-1 cue, **B.** for the phase I 1000ms Neutral-2 cue, **C.** for the phase II 1000ms Neutral cue, and **D.** for the phase II 1000ms Win cue. Colored lines and points denote individual patient associations. Dotted black line denotes mean association. Associations based on linear regression models. Significance based on p < 0.05*.

## Discussion

This study investigated the role of human dopamine concentrations in reproducing intervals of time from predictable cues in the presence and absence of reinforcement. We observed significant differences in the reproduction of time intervals and the generation of temporal errors between neurologically healthy individuals and patients with Parkinson’s disease. To understand how dopamine may be affecting these temporal learning measures, we utilized patients with Parkinson’s disease to directly measure real-time dopamine concentrations from the human striatum. We identified significant associations between human dopamine concentrations and the generation of temporal errors, which may be driving temporal learning behaviors. Specifically, we found associations between dopamine concentrations and temporal learning in the presence, but not in the absence, of reinforcement. These findings indicate that dopamine may be driving human temporal learning from reinforcements, and they may serve as a step to better understanding how humans perceive time, in general.

Using a peak interval procedure (**Fig. 1**), we measured the reproduction of 1000ms, 3000ms, and 5000ms intervals of time in both patients with Parkinson’s disease and neurologically healthy controls. Patients with Parkinson’s disease, who experience the progressive loss of midbrain dopamine neurons (Dauer & Przedborski, 2003), exhibited significant differences in both interval reproduction (**Fig. 2**) and temporal learning (**Fig. 3**) of 1000ms durations compared to neurologically healthy controls. Specifically, patients with Parkinson’s disease tended to overestimate 1000ms durations, whereas healthy subjects tended to underestimate 1000ms durations. This is consistent with other timing alterations found in patients with Parkinson’s disease, as decreased dopamine availability has been associated with a reduced ability to discriminate between time durations and an increased reporting of intervals as “long” (Parker et al. 2013; DiMarco et al. 2023; Sadibolova et al. 2024).

Interestingly, when measuring differences between patients with Parkinson’s disease and healthy controls, we did not find significant differences in the reproduction of 3000ms and 5000ms intervals of time. This finding aligns with other literature that has described nuanced differences in the reproduction and discrimination of intervals that fall between 500ms and 1200ms in duration and those greater than 1200ms in duration (Artieda et al. 1992; Koch et al. 2008; DiMarco et al. 2023). Specifically, patients with Parkinson’s disease have demonstrated a decreased ability to discriminate between intervals of time between 400ms and 600ms in duration, but not between intervals of greater than 2200ms (Koch et al. 2008). Patients with Parkinson’s disease have also presented with variable effects of prescribed dopaminergic therapies when performing timing tasks on intervals of less than 1000ms (Artieda et al. 1992; DiMarco et al. 2023). Therefore, it has been hypothesized that different circuitry may be involved in the perception of durations of time less than and greater than approximately 1200ms (Rammsayer & Troche, 2014). Dopaminergic action in the striatum has been reported to play a role in the timing of intervals greater than approximately 350ms (Sadibolova et al. 2023), whereas the prefrontal cortex and other “decision-making centers” have been shown to be engaged when timing intervals greater than about 1200ms, as timing longer intervals has been thought to require working memory and temporal planning processes (Rubia et al. 1998; Smith & Jonides 1999). Our results provide additional evidence of a neurochemical and behavioral difference between timing intervals 1000ms in duration and intervals of 3000ms and 5000ms in duration. These findings specifically may indicate that midbrain dopamine availability is integral in the reproduction of 1000ms durations.

Therefore, we focused the rest of our analyses on the 1000ms cues. Our results show that absolute temporal error decreases over repeated presentation of 1000ms cues of the reinforcement same type, demonstrating temporal learning (**Fig. 3**). As this form of task utilizes instrumental conditioning to teach subjects intervals of time, we sought to determine the role that real-time dopamine concentrations from the human striatum play in temporal learning behaviors. To investigate, we directly measured dopamine from patients with Parkinson’s disease using human voltammetry. We found that temporal learning occurred most for reinforced cues, as absolute temporal error decreased over repeated appearances for the 1000ms Win cue type (**Fig. 3**). We also found that increased absolute temporal error was associated with increased dopamine concentrations. Specifically, absolute temporal error was positively associated with dopamine concentrations for the reinforced cues, but it was not associated with absolute temporal error for the non-reinforced cues (**Fig. 4**). This provides evidence that dopamine may be acting as a teaching signal in response to reinforcement, which initially interferes with learning by increasing temporal error. We then showed that dopamine concentrations decreased across repeated presentation of reinforced cues, but again, not unreinforced cues (**Fig. 5**). Therefore, as a temporal cue is learned from reinforcement, dopamine decreases as temporal error also decreases.

It has been consistently reported that dopamine modulates timing behaviors (Meck 1996; Soares et al. 2016; Sadibolova et al. 2024), but it is still unknown how dopamine interacts with task structure and reinforcements to produce the human perception of time. Theories have posited that dopamine may be interacting locally through mechanisms related to reward prediction error signaling to drive population level activity (Paton & Buonomano, 2018). These population clock models would imply that local striatal dopamine dynamics may drive changes in interval timing by increasing dopamine concentrations leading to the under-reproduction (or underestimation) of intervals (Mello et al. 2015; Monteiro et al. 2023). As we found that patients with Parkinson’s disease on-average tend to over-reproduce (or overestimate) intervals (**Fig. 2**), it is possible that the increase in dopamine that is driving temporal learning in these patients is also linked to increased underestimation of intervals as patients learn overtime and exhibit fewer temporal errors. This would provide support for the population clock hypothesis of interval timing as increased striatal dopamine would theoretically result in a deceleration in population clock dynamics and temporal under-reproduction (underestimation) (Mello et al. 2015; Paton & Buonomano, 2018; Monteiro et al. 2023). As our sample is relatively small, more research is necessary to understand tonic and phasic dopamine changes, and how they may influence alterations in temporal behaviors during timing tasks with and without reinforcements.

Previous research has shown that on interval timing tasks, humans may not be aware of the direction of their temporal errors (Badar & Wiener, 2021; Badar & Weiner 2024; DiMarco et al. 2024). Specifically, Bader and Wiener et al. 2021 showed that subjects performing a reproduction task with non-directional feedback may not have been consciously aware of the direction of their temporal errors (over- or under-reproducing time intervals). Our results support this finding, as we also presented subjects with non-directional monetary feedback, and they exhibited improved reproduction over repeated cue presentation (**Fig. 3**). Our work extends these behavioral results by providing a potential explanation for the lack of awareness of the direction of temporal errors. We show that dopamine may be acting as a teaching signal, increasing in response to absolute temporal error (non-directional error) to correct timing behavior on subsequent presentation. Therefore, dopamine acting without directionality to modulate interval timing eliminates the need for human awareness of the directionality of temporal error signaling, instead only necessitating awareness if a temporal error occurs. This may specifically be related to the expectations surrounding the presented cues. As subjects may form an expected value of the temporal cue based on monetary reinforcement (DiMarco et al. 2024), it is possible that changes in this expected value (non-directional changes) are what drives this internal temporal error representation.

We found that increased temporal error is related to increased dopamine concentrations in the presence of reinforcements (**Fig. 4**). Therefore, the neurochemistry and behavior yielded from tasks that utilize instrumental conditioning to train subjects, or model organisms, to correctly reproduce time intervals may be driven by this temporal learning process. Although studies have shown that in model organisms, dopamine may modulate time independently of its modulation of reward processing (Soares et al. 2016). Most model organisms need to be highly trained with predictable cues and reinforcers to complete a timing task. Therefore, this reported dopaminergic signaling may be related to the generation of temporal errors in the presence of reinforcers. More work is needed to determine exactly how time perception relates to reinforcement processing, but this study provides evidence that tasks designed to measure timing may be measuring temporal learning behaviors, rather than strictly time perception, which we show are driven by dopamine. Therefore, future work should incorporate theories of reinforcement learning – specifically temporal difference reinforcement learning – to determine if time perception can be explained with dopaminergic signaling related to reward prediction error signaling.

This study examined the role of real-time human dopamine concentrations in concurrently collected timing behavior. It has implications in how we understand how task structure may relate to measured timing behaviors, as temporal learning may be occurring during interval timing tasks that utilize instrumental conditioning. We also provide evidence that dopamine may be driving behaviors related to temporal learning, particularly in response to reinforcements. Therefore, future work would need to consider the intersection of training, predictable cues, and reinforcement context when measuring timing behavior to understand how humans perceive and process time.

## Acknowledgments

We would like to thank our volunteers for their participation.

## References

1. Artieda, J., Pastor, M. A., Lacruz, F., & Obeso, J. A. (1992). Temporal discrimination is abnormal in Parkinson’s disease. Brain: a journal of neurology, 115 Pt 1, 199–210. 10.1093/brain/115.1.199

2. Bader F, Wiener M (2024). Neuroimaging Signatures of Metacognitive Improvement in Sensorimotor Timing. Journal of Neuroscience 28 February 2024, 44 (9) e1789222023; 10.1523/JNEUROSCI.1789-22.2023

3. Bang, D., Kishida, K. T., Lohrenz, T., White, J. P., Laxton, A. W., Tatter, S. B., Fleming, S. M., & Montague, P. R. (2020). Sub-second Dopamine and Serotonin Signaling in Human Striatum during Perceptual Decision-Making. Neuron, 108(5), 999–1010.e6. 10.1016/j.neuron.2020.09.015

4. Dauer, W., & Przedborski, S. (2003). Parkinson’s disease: mechanisms and models. Neuron, 39(6), 889–909. 10.1016/s0896-6273(03)00568-3

5. DiMarco, E.K., Shipp, A.R., & Kishida, K.T. (2024). Expected reward value and reward prediction errors reinforce but also interfere with human time perception. bioRxiv 2024.04.17.589985. 10.1101/2024.04.17.589985

6. DiMarco, E. K., Sadibolova, R., Jiang, A., Liebenow, B., Jones, R. E., Haq, I. U., Siddiqui, M. S., Terhune, D. B., & Kishida, K. T. (2023). Time perception reflects individual differences in motor and non-motor symptoms of Parkinson’s disease. Parkinsonism & related disorders, 114, 105800. 10.1016/j.parkreldis.2023.105800

7. Fung, B. J., Sutlief, E., & Hussain Shuler, M. G. (2021). Dopamine and the interdependency of time perception and reward. Neuroscience and biobehavioral reviews, 125, 380–391. 10.1016/j.neubiorev.2021.02.030

8. Hollerman, J. R., & Schultz, W. (1998). Dopamine neurons report an error in the temporal prediction of reward during learning. Nature neuroscience, 1(4), 304–309. 10.1038/1124

9. Hinton, S. C., & Meck, W. H. (2004). Frontal-striatal circuitry activated by human peak-interval timing in the supra-seconds range. Brain research. Cognitive brain research, 21(2), 171–182. 10.1016/j.cogbrainres.2004.08.005

10. Kishida, K. & Sands, L.P. (2021) A Dynamic Affective Core to Bind the Contents, Context, and Value of Conscious Experience. Affect Dynamics. ISBN : 978-3-030-82964-3

11. Kishida, K. T., Saez, I., Lohrenz, T., Witcher, M. R., Laxton, A. W., Tatter, S. B., White, J. P., Ellis, T. L., Phillips, P. E., & Montague, P. R. (2016). Subsecond dopamine fluctuations in human striatum encode superposed error signals about actual and counterfactual reward. Proceedings of the National Academy of Sciences of the United States of America, 113(1), 200–205. 10.1073/pnas.1513619112

12. Kishida KT, Sandberg SG, Lohrenz T, Comair YG, Sáez I, Phillips PEM, et al. (2011) Sub-Second Dopamine Detection in Human Striatum. PLoS ONE 6(8): e23291. 10.1371/journal.pone.0023291

13. Koch, G., Costa, A., Brusa, L., Peppe, A., Gatto, I., Torriero, S., Gerfo, E. L., Salerno, S., Oliveri, M., Carlesimo, G. A., & Caltagirone, C. (2008). Impaired reproduction of second but not millisecond time intervals in Parkinson’s disease. Neuropsychologia, 46(5), 1305–1313. 10.1016/j.neuropsychologia.2007.12.005

14. Liebenow, B., Jiang, A., DiMarco, E., Wilson, T., Siddiqui, M. S., Ul Haq, I., Laxton, A. W., Tatter, S. B., & Kishida, K. T. (2023). Intracranial subsecond dopamine measurements during a “sure bet or gamble” decision-making task in patients with alcohol use disorder suggest diminished dopaminergic signals about relief. Neurosurgical focus, 54(2), E3. 10.3171/2022.11.FOCUS22614

15. MacInnis, M.L.M., & Guilhardi, P. (2006). Basic interval discrimination procedures. In M. A. Anderson (Ed.), Tasks and Techniques: A Sampling of Methodologies for the Investigation of Animal Learning, Behavior, and Cognition, pp. 233–244. Hauppauge, NY: Nova Science Publishers. SBN 1-60021-126-7.

16. Meck WH. Internal clock and reward pathways share physiologically similar information-processing stages. Quant Anal Behav Biol Determ Reinf. 2014;7:121–138.

17. Meck W. H. (2005). Neuropsychology of timing and time perception. Brain and cognition, 58(1), 1–8. 10.1016/j.bandc.2004.09.004

18. Meck W. H. (1996). Neuropharmacology of timing and time perception. Brain research. Cognitive brain research, 3(3-4), 227–242. 10.1016/0926-6410(96)00009-2

19. Mello GBM, Soares S, Paton JJ (2015). A scalable population code for time in the striatum. Curr. Biol. 25, 1113–1122.

20. Montague, P. R., & Kishida, K. T. (2018). Computational Underpinnings of Neuromodulation in Humans. Cold Spring Harbor symposia on quantitative biology, 83, 71–82. 10.1101/sqb.2018.83.038166

21. Monteiro, T., Rodrigues, F. S., Pexirra, M., Cruz, B. F., Gonçalves, A. I., Rueda-Orozco, P. E., & Paton, J. J. (2023). Using temperature to analyze the neural basis of a time-based decision. Nature neuroscience, 26(8), 1407–1416. 10.1038/s41593-023-01378-5

22. Parker, K. L., Lamichhane, D., Caetano, M. S., & Narayanan, N. S. (2013). Executive dysfunction in Parkinson’s disease and timing deficits. Frontiers in integrative neuroscience, 7, 75. 10.3389/fnint.2013.00075

23. Paton, J. J., & Buonomano, D. V. (2018). The Neural Basis of Timing: Distributed Mechanisms for Diverse Functions. Neuron, 98(4), 687–705. 10.1016/j.neuron.2018.03.045

24. Qian, J., Hastie, T., Friedman, J., Tibshirani, R. and Simon, N. (2013). Glmnet for Matlab. http://hastie.su.domains/glmnet_matlab/

25. R Studio Team (2021). RStudio: Integrated Development for R. RStudio, PBC, Boston, MA URL http://www.rstudio.com/.

26. Rakitin, B. C., Gibbon, J., Penney, T. B., Malapani, C., Hinton, S. C., & Meck, W. H. (1998). Scalar expectancy theory and peak-interval timing in humans. Journal of experimental psychology. Animal behavior processes, 24(1), 15–33. 10.1037//0097-7403.24.1.15

27. Rammsayer, T. H., & Troche, S. J. (2014). In search of the internal structure of the processes underlying interval timing in the sub-second and the second range: A confirmatory factor analysis approach. Acta Psychologica, 147, 68–74. 10.1016/j.actpsy.2013.05.004

28. Rubia, K., Overmeyer, S., Taylor, E., Brammer, M., Williams, S., Simmons, A., Andrew, C., & Bullmore, E. (1998). Prefrontal involvement in “temporal bridging” and timing movement. Neuropsychologia, 36(12), 1283–1293. 10.1016/s0028-3932(98)00038-4

29. Sadibolova, R., Widmer, C., Fletcher, Z., Weill, S., & Terhune, D. B. (2023, April 17). Uncovering the latent structure of human time perception. 10.31234/osf.io/b92hn

30. Sadibolova, R., DiMarco, E.K., Jiang, A., Maas, B., Tatter, S.B., Laxton, A., Terhune, D.B., Kishida, K.T. (2024). Sub-second and multi-second dopamine dynamics underlie variability in human time perception. medRxiv 2024.02.09.24302276. doi: 10.1101/2024.02.09.24302276

31. Sands, L. P., Jiang, A., Liebenow, B., DiMarco, E., Laxton, A. W., Tatter, S. B., Montague, P. R., & Kishida, K. T. (2023). Subsecond fluctuations in extracellular dopamine encode reward and punishment prediction errors in humans. Science advances, 9(48), eadi4927. 10.1126/sciadv.adi4927

32. Smith, E. E., & Jonides, J. (1999). Storage and executive processes in the frontal lobes. Science (New York, N.Y.), 283(5408), 1657–1661. 10.1126/science.283.5408.1657

33. Soares, S., Atallah, B. V., & Paton, J. J. (2016). Midbrain dopamine neurons control judgment of time. Science (New York, N.Y.), 354(6317), 1273–1277. 10.1126/science.aah5234

34. The MathWorks Inc. (2022). MATLAB version: 9.13.0 (R2022b), Natick, Massachusetts: The MathWorks Inc. https://www.mathworks.com.

